# Bidirectional Modulation of Somatostatin-expressing Interneurons in the Basolateral Amygdala Reduces Neuropathic Pain Perception in Mice

**DOI:** 10.1101/2025.03.28.645947

**Authors:** Aditya Apte, Julia Fernald, Cody Slater, Marc Sorrentino, Brett Youngerman, Qi Wang

## Abstract

Neuropathic pain is characterized by mechanical allodynia and thermal (heat and cold) hypersensitivity, yet the underlying neural mechanisms remain poorly understood. This study examines the role of inhibitory interneurons in the basolateral amygdala (BLA) in modulating pain perception following nerve injury. Chemogenetic excitation of parvalbumin-positive (PV^+^) interneurons significantly alleviated mechanical allodynia but had minimal effects on thermal hypersensitivity. However, inhibition of PV^+^ interneurons did not produce significant changes in pain sensitivity, suggesting that reductions in perisomatic inhibition do not contribute to chronic pain states. In contrast, bidirectional modulation of somatostatin-positive (SST^+^) interneurons influenced pain perception in a modality-specific manner. Both excitation and inhibition of SST^+^ interneurons alleviated mechanical allodynia, indicating a potential compensatory role in nociceptive processing. Additionally, SST^+^ neuron excitation reduced cold hypersensitivity without affecting heat hypersensitivity, whereas inhibition improved heat hypersensitivity but not cold responses. These findings suggest that, in addition to PV^+^ neurons, SST^+^ interneurons in the BLA play a complex role in modulating neuropathic pain following nerve injury and may serve as a potential target for future neuromodulation interventions in chronic pain management.

## Introduction

Neuropathic pain is a chronic and debilitating condition that arises from damage to the nervous system, often manifesting in the absence of noxious stimuli. This form of pain is typically associated with disorders such as diabetes, trauma, or neurodegenerative diseases, and is characterized by spontaneous bouts of pain, allodynia, and hyperalgesia. The global prevalence of neuropathic pain is estimated at approximately 8% of the population and has a substantial impact on quality of life, making it a significant public health issue (Bennett and Xie, 1988; Yu et al., 2020). In addition to its negative impact on the quality of life, recent studies have suggested that severe chronic pain is strongly correlated with the rate of cognitive decline and an elevated risk of dementia (Zhao et al., 2023). Despite the burden of neuropathic pain to society, many current pharmacological approaches fail to provide sustained relief without undesirable side effects (Finnerup et al., 2015). Consequently, there is a critical need for better therapeutic interventions that target the underlying neural mechanisms of pain perception (Baron et al., 2010).

Perceptual processing is a highly adaptive process that depends on the characteristics of sensory inputs and behavioral state (Benucci et al., 2013; Yang and O’Connor, 2014; Zheng et al., 2015; Delis et al., 2018; Rodenkirch et al., 2019). Chronic pain is increasingly recognized as a disorder of maladaptive neural plasticity, involving long-term changes in synaptic strength, inhibitory tone, and neuromodulatory systems (Shiers et al., 2018; Guo et al., 2022). The basolateral amygdala (BLA) is a key structure in this process, integrating sensory input with emotional valence and contributing to the persistence of pain states (Neugebauer et al., 2004; Thompson and Neugebauer, 2017; Neugebauer, 2020). Lesion studies and electrophysiological recordings have demonstrated that neuronal activity within the BLA correlates with both the intensity and affective dimensions of pain (Ji and Neugebauer, 2009), with evidence suggesting that synaptic and cellular reorganization within this region facilitates the transition from acute to chronic pain (Kiritoshi et al., 2024).

A growing body of work suggests that inhibitory interneurons in the BLA play a pivotal role in regulating pain-related plasticity. Parvalbumin-expressing (PV^+^) interneurons provide perisomatic inhibition that stabilizes excitatory network activity and contributes to pain modulation (Caillard et al., 2000). Conversely, somatostatin-expressing (SST^+^) interneurons primarily target distal dendrites, influencing synaptic integration and network dynamics (Lovett-Barron et al., 2012; Urban-Ciecko and Barth, 2016). While PV^+^ and SST^+^ interneurons are well-characterized in cortical circuits, their functional contributions to BLA-dependent pain modulation remain incompletely understood.

Emerging research shows that these interneuron populations exhibit anatomical and functional heterogeneity. Previous studies have identified multiple subgroups of SST^+^ interneurons in the central nucleus of the amygdala and BLA, some of which project to downstream autonomic and nociceptive processing centers (McDonald and Mascagni, 2002; Muller et al., 2007; McDonald and Zaric, 2015; Mihaljevic et al., 2019; Bartonjo and Lundy, 2020, 2022). Similarly, PV^+^ interneurons in the BLA have been implicated in the regulation of oscillatory network activity and sensory gating (Yau et al., 2021; Amaya et al., 2024). Given the involvement of these interneurons in regulating emotional and sensory processing, it is plausible that their dysfunction contributes to chronic pain pathology.

In this study, we investigated the specific contributions of PV^+^ and SST^+^ interneurons in regulating nociceptive processing in the BLA. Using a chemogenetic approach, we examined how manipulation of these interneuron subtypes affects mechanical, thermal, and cold allodynia. Our findings suggest that SST^+^ and PV^+^ neurons exert distinct yet complementary roles in shaping pain-related behaviors, with potential implications for targeted neuromodulation therapies.

## Methods

### Animals and Surgical Procedures

All experimental procedures were approved by the Columbia University Institutional Animal Care and Use Committee (IACUC) and were conducted with compliance with NIH guidelines. Adult mice of both male and female sex (12 female and 12 male), aged 3 ∼ 6 months, were used in the experiments. The strains used were SST-IRES-Cre (RRID: IMSR_JAX:013044) and PV-IRES-Cre (RRID: IMSR_JAX:017320). All mice were kept under a 12-hour light-dark cycle.

#### Intracranial adeno-associated virus injection

The procedure for injecting the viral vector solution was previously described in detail (Liu et al., 2024). Briefly, mice were anesthetized using isoflurane (5% for induction and 2% for maintenance) and placed in a stereotaxic frame (Kopf Instruments, Tujunga, CA). Body temperature was maintained at 38 °C, using a feedback-controlled heating pad (FHC, Bowdoinham, ME). Once the animal’s condition stabilized, Buprenorphine (0.05 mg/kg) was administered subcutaneously. During aseptic preparation, fur on the skull of the mouse was shaved clean, creating a site for incision. Next, three alternating passes of alcohol swabs and betadine were used to sterilize the site, followed by subcutaneous injection of lidocaine (2%). After exposing the skull, a burr hole was drilled above the right BLA (AP: -1.4 mm, ML: 2.6 mm, DV: -4.9 mm), with saline applied to the craniotomy to prevent drying of the brain surface. We targeted the right BLA based on evidence from earlier research that linked tactile hypersensitivity to increased activity in the amygdala on the right side of the mouse’s brain (Carrasquillo and Gereau, 2008; Ji and Neugebauer, 2009; Kolber et al., 2010; Li et al., 2017). Pulled capillary glass micropipettes were back-filled with 150 nL AAV solution (pAAV1-hSyn-DIO-hM3D(Gq)-mCherry or pAAV1-hSyn-DIO-hM4D(Gi)-mCherry), which was subsequently injected into the BLA at 0.7nL/s using a precision injection system (Nanoliter 2020, World Precision Instruments, Sarasota, FL). The pipette was left in place for at least 10 minutes following injection and slowly withdrawn. Following withdrawal of the pipette, skin was closed with Vetbond. Baytril (5 mg/kg) and Ketoprofen (5 mg/kg) were administered postoperatively for 5 days.

#### Constriction nerve injury (CNI) procedure

Nerve injury surgeries were performed 10 days following AAV injection surgery. The injury was induced using a chronic constriction method similar to previously described (Austin et al., 2012). In aseptic surgeries, mice were initially anesthetized with isoflurane. To access the left lateral thigh of the mouse, the animal was placed on its side, and limbs were secured to the surgical surface to ensure stability of the leg during surgery. A 1 cm thick gauze pad was placed between the hind legs to provide support to the leg and make access more feasible. After shaving the surgical site, a 1 cm long incision was made in the proximal one third of the lateral thigh of the left leg. The sciatic nerve was exposed by opening muscle fascia between the gluteus superficialis and biceps femoris muscles using sterilized toothpicks to push apart muscle fibers without causing damage to nearby blood vessels. The sciatic nerve was then raised from the cavity using fine-tip forceps inserted under the nerve and gently stretched using said forceps. Three nylon ligatures are tied around the sciatic nerve 2 mm apart, to cause enough constriction to the nerve without preventing epineural blood flow. The wound was closed with absorbable sutures in the muscle and skin. Baytril (5 mg/kg) was administered subcutaneously and Triple antibiotics ointment was applied for three days in post-operative care. The animal is allowed to recover from surgery for 3 days before injured baseline behavioral testing begins.

### Chemogenetic Manipulation

To test the role of SST+ or PV+ interneurons in the BLA in pain behavior, we performed 7 days of chemogenetic manipulation (i.e., treatment) and 3 days of saline control (sham-treatment). For all chemogenetic manipulation experiments, clozapine-N-oxide (CNO; Hello Bio, NJ) was injected i.p. (3 mg/kg body weight) 30 minutes prior to testing. On sham-treatment days, an equivalent volume of saline was injected i.p. 30 minutes prior to testing. Treatment and sham-treatment sessions were randomly interleaved and were blind to the experimenter who conducted behavioral tests.

### Nociceptive Behavioral Assays

All behavioral assays were performed by an experimenter blinded to the mouse group and administered treatment. 5 sessions of baseline data were collected prior to surgery. 5 sessions of injured baseline data were collected following CNI, and 10 days of data collection in response to treatment/sham-treatment was performed following conclusion of injured baseline data collection. (**Figure 1A**)

**Figure 1.**
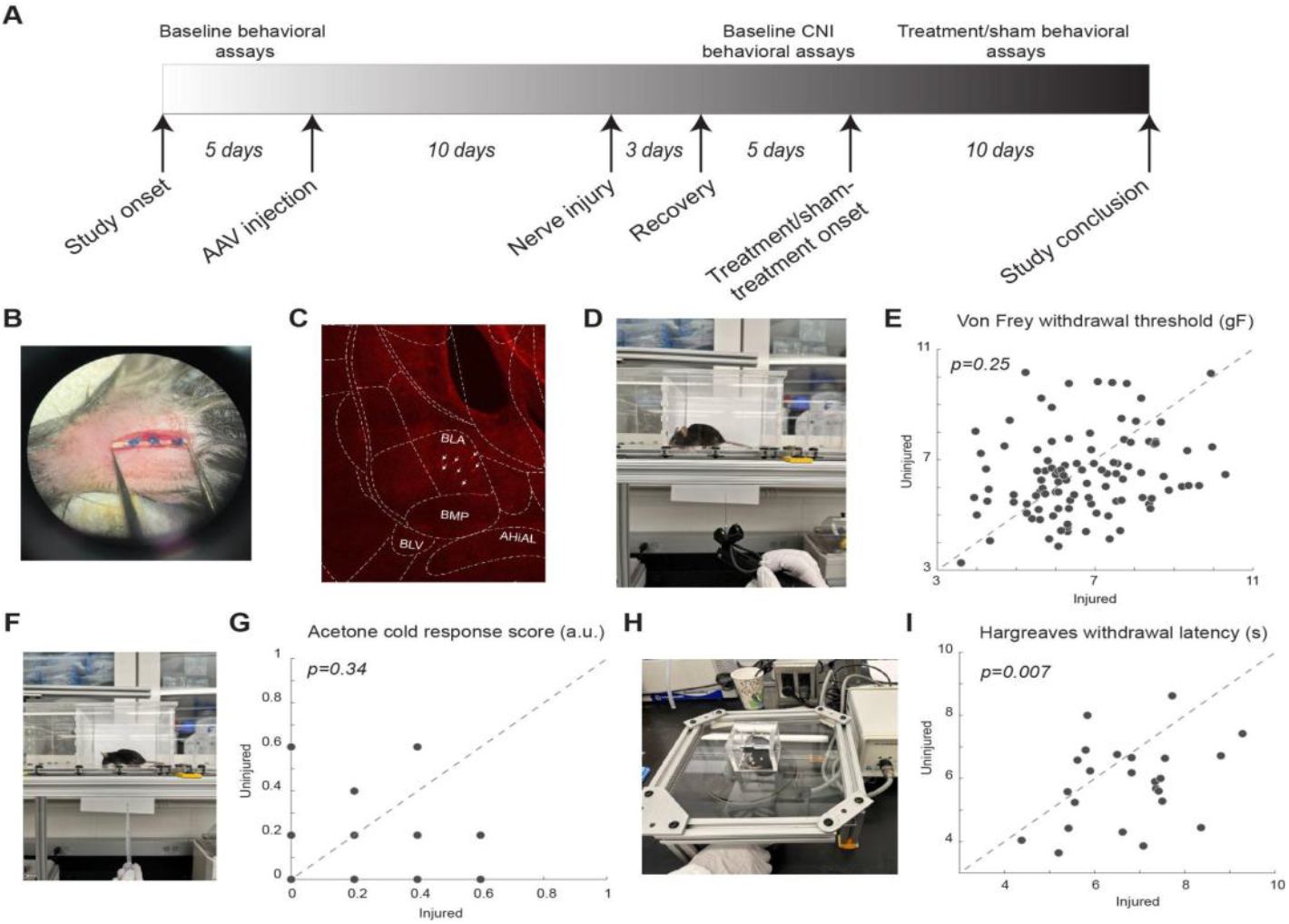
Experimental setup and baseline results of the behavioral assays. **A)** The timeline of the study. **B)** Photo of example CNI. **C)** Example immunohistology confirming expression of DREADD receptors in SST+ neurons in the BLA. White arrows indicate SST+ neurons. **D**,**E)** Photo of a von Frey test and withdrawal thresholds in von Frey tests prior to CNI. **F**,**G)** Photo of an acetone test and response scores in acetone tests prior to CNI. **H**,**I)** Photo of a Hargreaves test and withdrawal latencies in Hargreaves tests prior to CNI.

#### Von Frey tests for mechanical allodynia

Mice were habituated on an elevated acrylic mesh platform in 4 × 4 × 3 inch acrylic boxes for thirty minutes before mechanical stimulation with a von Frey filament (**Figure 1B**) (Wilson et al., 2019). The sensitivity of the paw to mechanical stimulation was measured using an electronic von Frey aesthesiometer (eVF, Model: 38450, Ugo Basile). Once the filament is applied, the eVF records the force readout (gF) and latency (s). Mechanical paw withdrawal threshold was defined as the force applied by the filament causing the paw to withdraw. Three measurements were taken for each of the hindpaws, ipsilateral and contralateral to the injury side, per session.

#### Hargreaves tests for heat sensitivity

Mice were habituated on an elevated ⅛ inch thick glass platform in a 2 × 2 × 2 inch ventilated acrylic box for thirty minutes prior to application of a thermal stimulus using a constant radiant heat source (Model: 7371, Ugo Basile) with an active intensity of 25% to the plantar surface of each hind paw through the glass surface (**Figure 1D**) (Wilson et al., 2019). The latency for paw withdrawal was measured five times per hindpaw. Three sessions of Hargreaves latency test were performed in each of the baseline, injured baseline, and treatment, with two sessions performed for sham-treatment.

#### Acetone test for cold sensitivity

Mice were habituated on an elevated acrylic mesh platform in 4 × 4 × 3 in acrylic boxes for thirty minutes prior to application of acetone droplet (**Figure 1E**). Acetone was loaded into a 5 mL pipette and a drop was lightly applied through the acrylic mesh to the plantar surface of the hindpaw, without touching the paw with the pipette to avoid false response (Wilson et al., 2019). Nociceptive responses after applying acetone droplet to the mouse’s hindpaw were quantified using a version of the scoring system described (Colburn et al., 2007) where 0 = minimal to no lifting, licking, or shaking of the hindpaw; 1 = lifting, licking, and/or shaking of the hindpaw, continuing for between 1-5 s; 2 = lifting, licking, and/or shaking of the hindpaw, prolonged or repetitive, extending beyond 5s after application of acetone (Colburn et al., 2007). Responses to acetone were gauged 5 times per hindpaw per session. Three sessions of the acetone evaporation test were performed in each of the baseline, injured baseline, and treatment, with two sessions performed for sham-treatment.

### Immunohistochemistry

IHC procedures to confirm the expression of DREADD receptors in the BLA were similar to those previously described (Liu et al., 2024). Briefly, at the end of the study, a subset of mice was anesthetized and transcardially perfused with phosphate buffered saline (PBS) followed immediately by ice-cold 4% paraformaldehyde (PFA). The brain was carefully extracted from the skull and post-fixed for twenty-four hours at 4 °C in 4% PFA, and then preserved in a 30% sucrose (wt/vol) PBS solution for 72 hours at 4 °C. After 3 days, extracted brains were embedded in optimum cutting temperature compound, and 30-μm coronal slices were obtained using a cryostat (Leica CM1950), and placed on a glass microscope slide. Brain slices were washed three times in PBS followed by placement and mounting of a coverslip using Fluoromount-G medium with DAPI. The slices were imaged using 20x under a confocal microscope (Nikon Ti2) with a spinning disk (Yokogawa CSU-W1).

### Data Analysis

All data analyses were conducted on individual animals. The averages and standard errors of means were then calculated across animals for each experimental group. For von Frey and Hargreaves tests, the normalized change between two hindpaws was calculated as 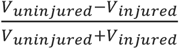, where *V*_*uninjured*_ is the measurement from the hindpaw on the uninjured side, while *V*_*injured*_ is the measurement from the hindpaw on the injured side. Because in acetone tests, the hindpaw on the injured side usually had a higher score, the normalized change was calculated as 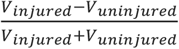, where *V*_*uninjured*_ is the measurement from the hindpaw on the uninjured side, while *V*_*injured*_ is the measurement from the hindpaw on the injured side. Following normalization, undefined values were excluded from the data analysis.

## Statistics

One-sample Kolmogorov-Smirnov test was used to verify the normality of the data. For data with a normal distribution, a Student’s t-test was performed. Otherwise, the Mann-Whitney U-test for unpaired samples or the Wilcoxon signed-rank test was used for paired samples. One-way ANOVA was performed to determine significance of normalized plots comparing injured baseline, activation/inhibition, and saline-control responses.

## Results

To assess the behavioral outcome of constriction nerve injury (CNI), we first measured the baseline sensitivity to mechanical (von Frey), thermal (infrared heat), and cold (acetone) stimuli prior to the induction of CNI (**Fig. 1**). Before the injury, we did not find significant differences between the two hindpaws in sensitivity to the mechanical (6.67 ± 0.14 vs 6.47 ± 0.14, p=0.25, paired Student’s t-test. **Fig. 1D**,**E**), and cold stimulus (0.175 ± 0.038 vs 0.125 ± 0.037, p=0.34, paired Student’s T-test. **Fig. 1F**,**G**). However, there was a small but significant difference in the latency in response to thermal stimuli between the two hindpaws (6.74 ± 0.25 vs 5.86 ± 0.27, p=0.007, paired Student’s T-test. **Fig. 1H**,**I**). After CNI, the mice exhibited hypersensitivity to mechanical and thermal stimulus in the injured hindpaw compared to the uninjured paw (**Fig. 2-4**), confirming the neuropathic pain model.

**Figure 2:**
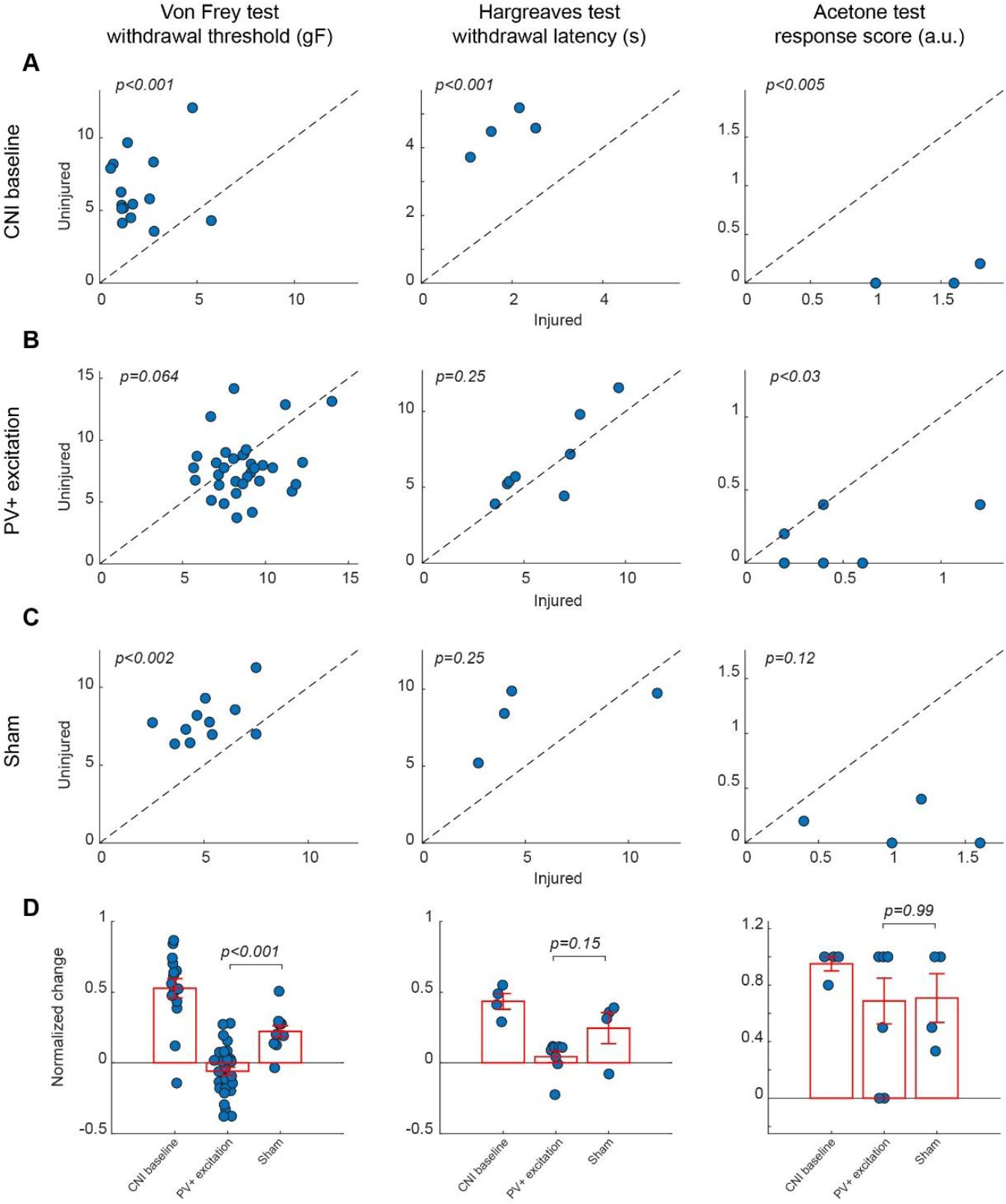
Effects of activation of PV+ interneuron in the BLA. **A)** Behavioral outcomes in the von Frey, Hargreaves, and acetone tests after CNI. **B)** Behavioral outcomes in the von Frey, Hargreaves, and acetone tests after CNI with activation of PV+ neurons in the BLA. **C)** Behavioral outcomes in the von Frey, Hargreaves, and acetone tests after CNI during sham control. **D)** Normalized differences between the injured and uninjured hindpaws during CNI baseline, excitation of PV+ neurons in the BLA, and sham control conditions.

### Excitation of PV+ interneurons in the BLA alleviates mechanical but not thermal allodynia

Following the CNI, mice developed robust mechanical, thermal, and cold hypersensitivity in the hindpaw on the injured side compared to the uninjured side. For mice with excitatory DREADD receptors expressed in parvalbumin-expressing (PV+) neurons in the basolateral amygdala (BLA), their mechanical paw withdrawal thresholds were significantly reduced for the injured side as compared to the uninjured side (2.03 ± 0.38 vs 6.38 ± 0.61 g, p<0.001, Wilcoxon Signed Rank Test, **Fig. 2A left panel**). Similarly, the withdrawal latencies of the paw on the injured side in response to infrared heat (Hargreaves test) were shorter (1.83 ± 0.32 vs 4.49 ± 0.29 s, p<0.001, Wilcoxon Signed Rank Test, **Fig. 2A middle panel**), and their acetone-evoked cold responses were enhanced compared to the hindpaw on the uninjured side (1.35 ± 0.21 vs 0.05 ± 0.05, p<0.005, paired Student’s T-test) (**Fig. 2A, right panel**). These changes were consistent with persistent allodynia and hyperalgesia induced by CNI (Bennett and Xie, 1988; Austin et al., 2012). Given that these animals showed little difference in these assays between the two paws prior to CNI, the observed hypersensitivity following nerve injury confirmed the replication of a neuropathic pain model upon which we will subsequently test the effects of various chemogenetic manipulations on pain perception.

Chemogenetic activation of PV-expressing GABAergic interneurons in the BLA significantly ameliorated neuropathic pain behaviors induced by CNI in von Frey tests. Following CNO-mediated activation of PV+ neurons, the mice exhibited a marked increase in mechanical withdrawal thresholds for the injured paw, resulting in a comparable threshold to the uninjured paw (8.69 ± 0.33 vs 7.85 ± 0.41, p=0.0644, Wilcoxon Signed Rank Test, **Fig. 2B**, left panel). As expected, the administration of saline, which served as a sham control for CNO-mediated activation of PV+ neurons in the BLA, did not decrease the difference in von Frey withdrawal threshold between the injured and uninjured hindpaws (5.13 ± 0.47 vs 7.9 ± 0.43, p<0.002, Wilcoxon, **Fig. 2C**, left panels).

To further quantify the effect of the excitation of PV+ neurons in the BLA, we calculated the normalized difference in all responses between the injured and uninjured paws, which is the difference of the measure between the two paws over the sum of the measure of the two paws (see Methods). A normalized difference of 0 would indicate that the injured and uninjured paws had the same response to the behavioral tests. For the von Frey test, the excitation of BLA PV+ neurons decreased the normalized difference induced by the CNI, and maintained a significant difference with response to saline (−0.059 ± 0.029 vs 0.22 ± 0.04, p<0.001, Mann-Whitney U-test, **Fig. 2D**, left panel).

Similarly, heat hypersensitivity was improved by the activation of PV+ neurons in the BLA, with differences in the withdrawal latencies between the two hindpaws vanishing after excitation of BLA PV+ neurons (6.03 ± 0.77 vs 6.65 ± 0.96, p=0.25, Wilcoxon Signed Rank Test, **Fig. 2B**, middle panel). However, in the saline control sessions, the difference between the two hindpaws was not statically significant (5.61 ± 1.96 vs 8.31 ± 1.08, p=0.25, Wilcoxon Signed Rank Test, **Fig. 2C**, middle panel), probably due to the small number of samples. When looking at the normalized difference between PV+ excitation and saline control sessions for Hargreaves tests, although the normalized difference during BLA PV+ excitation is much smaller than in sham control conditions, this difference was not statistically significant (0.04 ± 0.04 vs 0.24 ± 0.11, p=0.15, Mann-Whitney U-test, **Fig. 2D**, middle panel), suggesting that the excitation of PV+ interneurons in the BLA only minimally reduces thermal allodynia.

In the acetone induced cold test, we still observed a significant difference in the response score following the excitation of PV+ neurons in the BLA (0.475 ± 0.12 vs 0.125 ± 0.06, p<0.03, Wilcoxon Signed Rank Test, **Fig. 2B**, right panel). In the saline controls, the differences in the response score between the two hindpaws were not significant (1.05 ± 0.25 vs 0.15 ± 0.096, p=0.12, Wilcoxon Signed Rank Test, **Fig. 2C**, right panel). However, the normalized difference was about the same between the PV+ excitation sessions and the saline control sessions (0.68 ± 0.16 vs 0.71 ± 0.17, p=0.99, Mann-Whitney U-test, **Fig. 2D**, right panel), indicating that the excitation of PV+ interneurons in the BLA has no beneficial effects.

Taken together, these results suggest that the activation of PV+ interneurons in the BLA selectively alleviates mechanical allodynia while exerting little effects on heat and cold hypersensitivity.

### Inhibition of PV+ interneurons in the BLA had no effect on neuropathic hypersensitivity

In contrast, chemogenetic inhibition of PV+ interneurons in the BLA did not significantly improve mechanical, thermal, or cold hypersensitivity induced by CNI. In the von Frey tests, paw withdrawal thresholds differed significantly between ipsilateral and contralateral hindpaws following CNI (2.76 ± 0.39 vs 6.73 ± 0.45 g, p<0.001, Wilcoxon Signed Rank Test, **Fig. 3A**, left panel) and during the sham control (4.33 ± 0.52 vs 6.88 ± 0.57, p<0.001, Wilcoxon Signed Rank Test, **Fig. 3C**, left panel), indicating the injured side remained significantly more sensitive than the contralateral side. However, following CNO-mediated inhibition of PV+ interneurons, mechanical paw withdrawal thresholds retained a significant difference between the injured and uninjured sides (5.27 ± 0.31 vs 6.75 ± 0.31 g, p<0.0011, Wilcoxon Signed Rank Test, **Fig. 3B**, left panel). We next examined whether the normalized differences between the two hindpaws differed between the PV+ inhibition and sham control conditions. Although there was a decrease in normalized difference following inhibition of BLA PV+ neurons compared to sham control, the difference was marginally significant (0.25 ± 0.045 vs 0.13 ± 0.036, p=0.08, Mann-Whitney U-test, **Fig. 3D**, left panel), suggesting that the inhibition of PV+ neurons in the BLA had weak effects on mechanical hypersensitivity.

**Figure 3:**
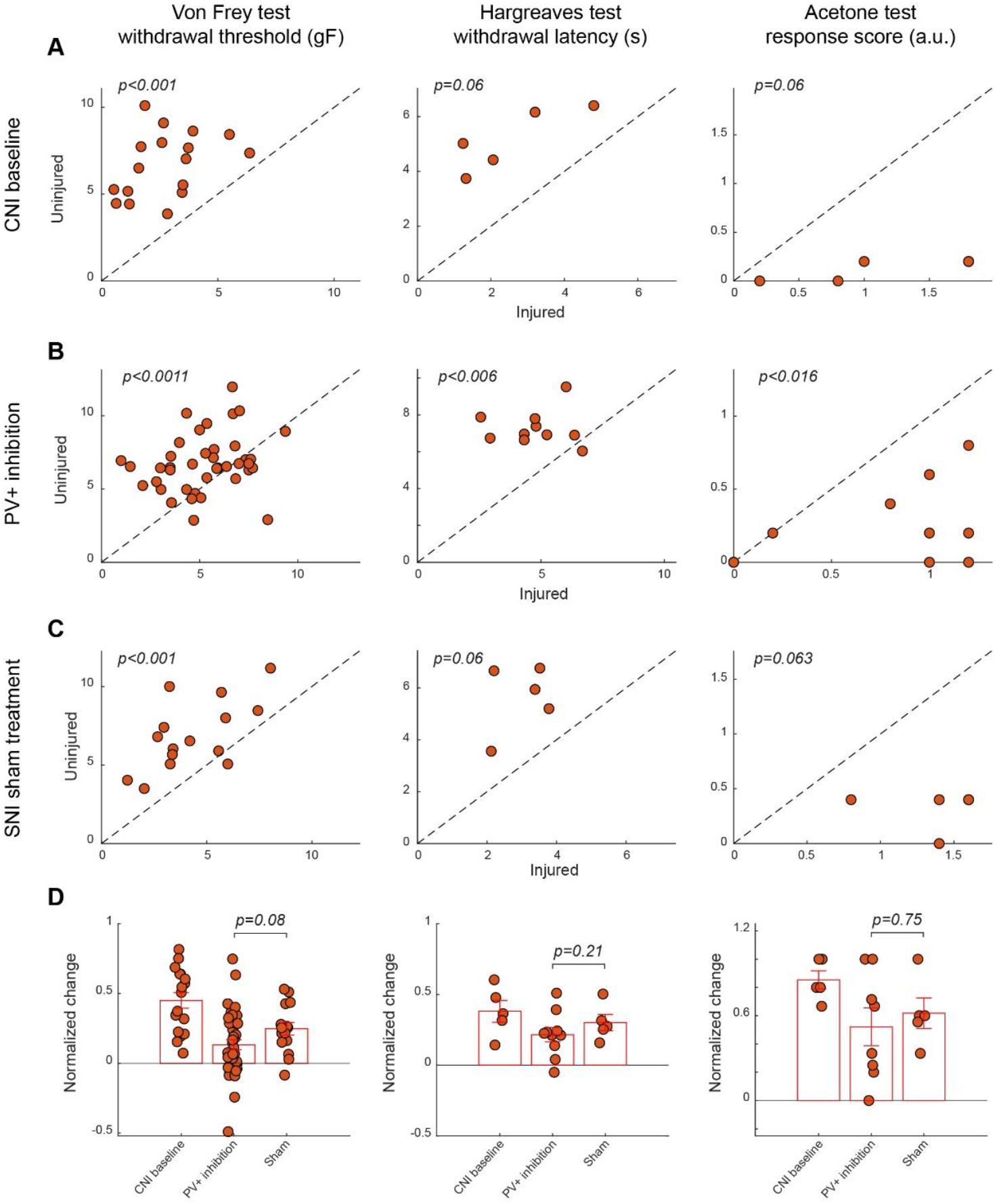
Effects of inhibition of PV+ interneuron in the BLA. **A)** Behavioral outcomes in the von Frey, Hargreaves, and acetone tests after CNI. **B)** Behavioral outcomes in the von Frey, Hargreaves, and acetone tests after CNI with inhibition of PV+ neurons in the BLA. **C)** Behavioral outcomes in the von Frey, Hargreaves, and acetone tests after CNI during sham control. **D)** Normalized differences between the injured and uninjured hindpaws during CNI baseline, inhibition of PV+ neurons in the BLA, and sham control conditions.

As with responses indicating mechanical paw withdrawal threshold, although the injured hindpaw exhibited hypersensitivity in Hargreaves test following CNI (2.52 ± 0.67 vs 5.15 ± 0.51, p=0.06, Wilcoxon Signed Rank Test, **Fig. 3A**, middle panel) and during sham control (3 ± 0.35 vs 5.86 ± 0.58, p=0.06, Wilcoxon Signed Rank Test, **Fig. 3C**, middle panel), the thermal sensitivity of the mice was not restored to near-contralateral latencies following inhibition of PV+ neurons in the BLA (4.8 ± 0.43 vs 7.3 ± 0.30, p<0.006, Wilcoxon Signed Rank Test, **Fig. 3B**, middle panel). Following calculation of normalized differences, we found that responses to PV+ inhibition and saline administration did not show a statistically significant difference (0.22 ± 0.05 vs 0.3 ± 0.06, p=0.21, Mann-Whitney U-test, **Fig. 3D**, middle panel), suggesting that inhibition of PV+ neurons in the BLA has no effect on heat allodynia.

Similarly, in the acetone induced cold test, although there was a significant difference between two hindpaws after CNI (1.12 ± 0.31 vs 0.12 ± 0.049, p=0.06, Wilcoxon Signed Rank Test, **Fig. 3A**, right panel), we still observed significant differences in the response following inhibition of BLA PV+ neurons (0.76 ± 0.16 vs 0.24 ± 0.09, p<0.016, Wilcoxon Signed Rank Test, **Fig. 3B**, right panel), while saline control results showed marginally significant difference between hindpaws (1.36 ± 0.15 vs 0.32 ± 0.08, p=0.063, Wilcoxon Signed Rank Test, **Fig. 3C**, right panel). Normalized difference analysis also confirmed that there is no significant difference in response between PV+ inhibition and saline control (0.52 ± 0.13 vs 0.61 ± 0.11, p=0.75, Mann-Whitney U-test, **Fig. 3D**, right panel). These findings align with previous reports (e.g., Corder et al., Science 2019) suggesting that inhibition of PV+ interneuron activity is insufficient to normalize sensory thresholds. The persistence of mechanical, thermal, and cold hypersensitivity under PV+ inhibition underscores the specificity of BLA PV+ neuron activation in counteracting allodynia.

### Excitation of SST+ interneurons in the BLA reduces mechanical and cold, but not heat hypersensitivity

After characterizing the effect of manipulation of PV+ neurons in the BLA on pain-related behavior, we next examined the role of somatostatin (SST)-expressing neurons in the BLA in pain perception. Activation of SST+ neurons in the BLA diminished mechanical allodynia induced by CNI (1.96 ± 0.29 vs 5.62 ± 0.26, p<0.001, Wilcoxon Signed Rank Test, **Fig. 4A**, left panel). Following CNO-mediated activation of BLA SST+ interneurons, mice showed a significant increase in mechanical paw-withdrawal threshold for the injured paw, reaching values nearly equivalent to uninjured paw responses (5.53 ± 0.31 vs 5.93 ± 0.27, p=0.1755, Wilcoxon Signed Rank Test, **Fig. 4B**, left panel). Consistent with results in PV+ manipulation experiments, sham control with saline administration did not decrease the difference in response between injured and uninjured paws (3.32 ± 0.37 vs 6.84 ± 0.55, p<0.001, Wilcoxon Signed Rank Test, **Fig. 4C**, left panel).

**Figure 4.**
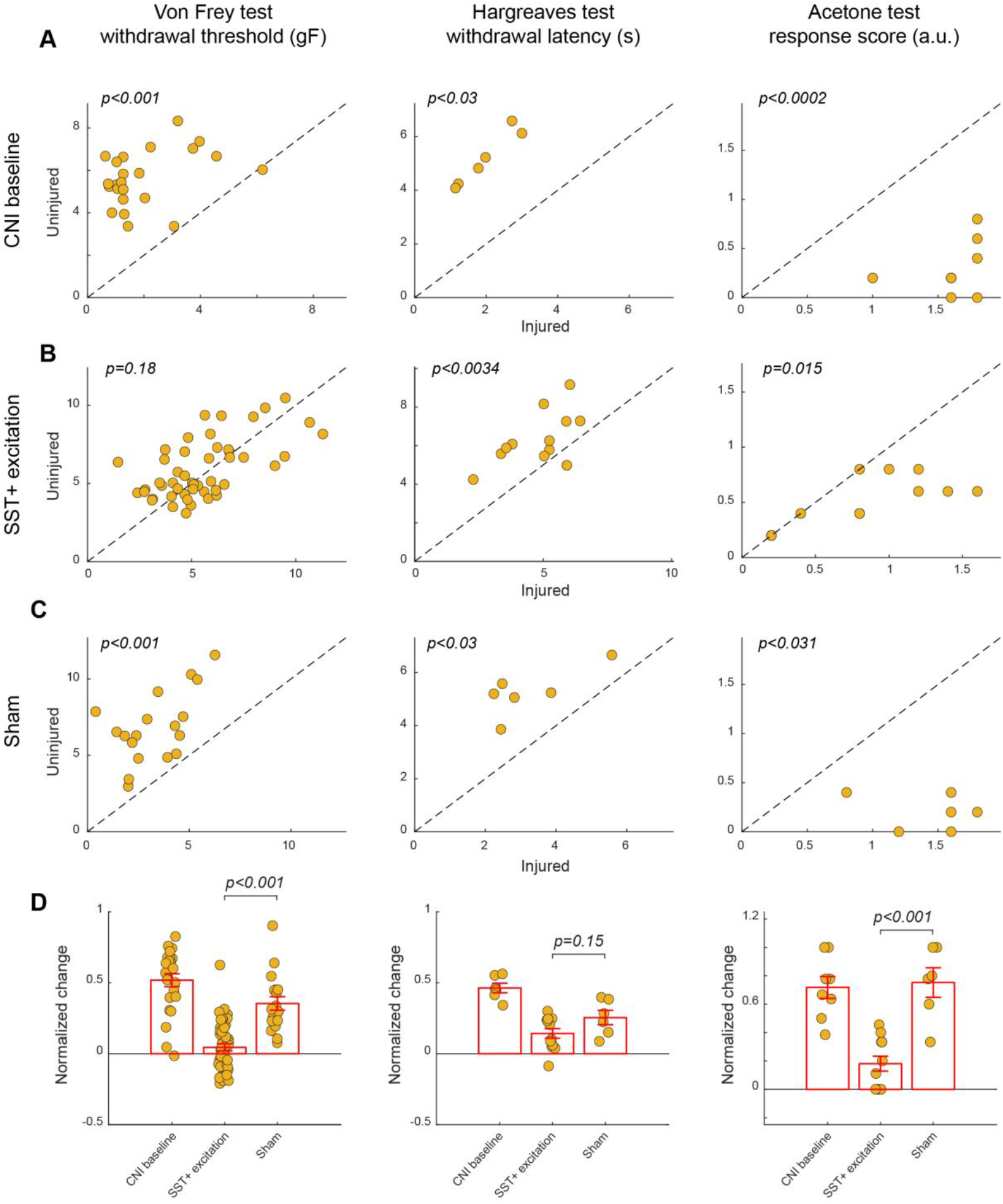
Effects of activation of SST+ interneuron in the BLA. **A)** Behavioral outcomes in the von Frey, Hargreaves, and acetone tests after CNI. **B)** Behavioral outcomes in the von Frey, Hargreaves, and acetone tests after CNI with activation of SST+ neurons in the BLA. **C)** Behavioral outcomes in the von Frey, Hargreaves, and acetone tests after CNI during sham control. **D)** Normalized differences between the injured and uninjured hindpaws during CNI baseline, excitation of SST+ neurons in the BLA, and sham control conditions.

We further calculated the normalized difference between the injured and uninjured paws for the von Frey test. Activation of BLA SST+ interneurons reduced the normalized difference caused by CNI. and retained a statistically significant difference with activation responses following normalization (0.35 ± 0.045 vs 0.045 ± 0.025, p<0.001, Mann-Whitney U-test, **Fig. 4D**, left panel).

Activation of BLA SST+ neurons failed to improve heat hypersensitivity as thermal withdrawal latencies differed significantly during CNI baseline (1.97 ± 0.31 vs 5.17 ± 0.41, p=0.03, Wilcoxon Signed Rank Test, **Fig. 4A**, middle panel), SST+ activation (4.82 ± 0.37 vs 6.34 ± 0.4, p<0.0034, Wilcoxon Signed Rank Test, **Fig. 4B**, middle panel), and sham control (3.23 ± 0.5 vs 5.27 ± 0.37, p=0.03, Wilcoxon Signed Rank Test, **Fig. 4C**, middle panel). There also was a non-significant difference in the normalized differences between SST+ activation and saline sham control (0.144 ± 0.03 vs 0.25 ± 0.051, p=0.15, Mann-Whitney U-test, **Fig. 4D**, middle panel). This suggests that SST+ activation does not reduce thermal allodynia.

Unlike the manipulation of PV+ neurons in the BLA, activation of SST+ neurons in the BLA appeared to improve cold-induced allodynia, as indicated by a reduction in acetone-evoked responses between injured and injured paws (0.81 ± 0.14 vs 0.5 ± 0.07, p<0.0156, Wilcoxon Signed Rank Test, **Fig. 4B**, right panel) as compared to in Saline control sessions (1.43 ± 0.15 vs 0.2 ± 0.07, p<0.031, Wilcoxon Signed Rank Test, **Fig. 4C**, right panel). This was further confirmed by their normalized differences (0.18 ± 0.05 vs 0.75 ± 0.1, p=0.001, Mann-Whitney, **Fig. 4D**, right panel.) These results highlight that SST+ interneuron excitation not only normalizes mechanical hypersensitivity, but also extends its analgesic effects to cold hypersensitivity.

### Inhibition of SST+ interneurons improves mechanical and heat, but not cold, hypersensitivity

We found that chemogenetic inhibition of SST+ neurons in the BLA also conferred a relief from mechanical and heat hypersensitivity. In response to CNO-mediated inhibition of SST+ neurons, there was a significant increase in mechanical paw withdrawal thresholds for the injured side to uninjured levels. (5.97 ± 0.25 vs 6.02 ± 0.27, p=0.88, Wilcoxon Signed Rank Test, **Fig. 5B**, left panel), while the significant difference between injured and uninjured hindpaws was observed in both CNI baseline (3.35 ± 0.2 vs 7.56 ± 0.34, p<0.001, Wilcoxon Signed Rank Test, **Fig. 5A**, left panel) and saline control sessions (3.66 ± 0.42 vs 6.56 ± 0.45, p<0.001, Wilcoxon Signed Rank Test, **Fig. 5C**, left panel). After calculation of normalized difference, there was a significant difference between BLA SST+ inhibition condition and saline control (−0.0022 ± 0.018 vs 0.302 ± 0.049, p<0.001, Mann-Whitney U-test, **Fig. 5D**, left panel).

**Figure 5:**
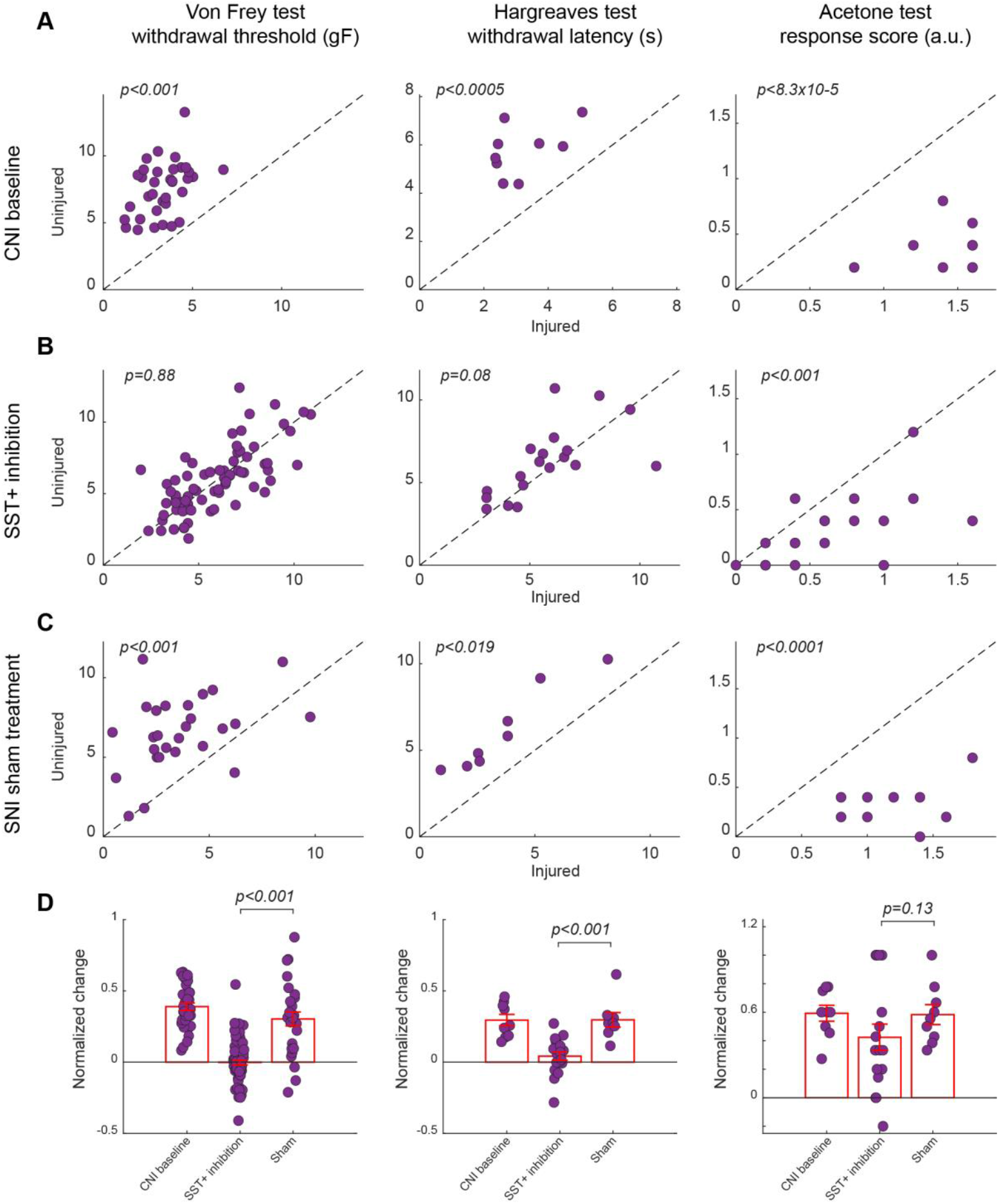
Effects of inhibition of SST+ interneuron in the BLA. **A)** Behavioral outcomes in the von Frey, Hargreaves, and acetone tests after CNI. **B)** Behavioral outcomes in the von Frey, Hargreaves, and acetone tests after CNI with inhibition of SST+ neurons in the BLA. **C)** Behavioral outcomes in the von Frey, Hargreaves, and acetone tests after CNI during sham control. **D)** Normalized differences between the injured and uninjured hindpaws during CNI baseline, inhibition of SST+ neurons in the BLA, and sham control conditions.

Unlike BLA SST+ activation, BLA SST+ inhibition resulted in improved thermal hypersensitivity, with injured paw response reaching contralateral levels (5.78 ± 0.48 vs 6.26 ± 0.49, p=0.08, Wilcoxon Signed Rank Test, **Fig. 5B**, middle panel). Responses to saline control mirrored CNI baseline response, maintaining a significant difference between injured and uninjured hindpaws (**Fig. 5A**, C, middle panel). There was also a significant difference between SST+ inhibition and saline administration when comparing the normalized difference (0.042 ± 0.028 vs 0.29 ± 0.051, p<0.001, Mann-Whitney U-test, **Fig. 5D**, middle panel).

However, the inhibition of BLA SST+ neurons did not affect the injured paw response to acetone-induced cold hypersensitivity (0.64 ± 0.1 vs 0.32 ± 0.07, p<0.001, Wilcoxon Signed Rank Test, **Fig. 5B**, right panel). There were also no statistically significant differences between CNI, SST+ inhibition, and saline administration when examining the normalized differences (p=0.13, Mann-Whitney U-test, **Fig. 5A, C, & D**, right panel).

## Discussion

Our results demonstrate that both PV^+^ and SST^+^ interneurons in the BLA contribute to the regulation of nociceptive processing, but they do so in modality-specific ways. Excitation of PV^+^ neurons in the BLA robustly alleviated mechanical allodynia. This is in line with the results of a previous study, where excitation of PV^+^ neurons in the spinal cord in mice with nerve injury attenuated their mechanical hypersensitivity, whereas transiently silencing the spinal cord PV+ neurons in naive mice resulted in mechanical allodynia (Petitjean et al., 2015). This improvement is likely because, in the BLA, perisomatic inhibition by PV^+^ neurons induces a strong and immediate decrease in the firing rate of excitatory neurons (Yau et al., 2021; Amaya et al., 2024). Upon nerve injury, the reduced expression of PV in PV^+^ neurons disrupted firing patterns, thereby decreasing the overall inhibition exerted by PV^+^ neurons (Qiu et al., 2024). Therefore, chemogenetic excitation of PV^+^ neurons restored inhibition exerted by PV^+^ neurons in the BLA and reinstated mechanical sensitivity. We failed to find PV^+^ inhibition to normalize hypersensitivity for mechanical, cold, or heat stimuli. This is consistent with previous studies suggesting that inhibition by PV^+^ neurons is already impaired following nerve injury (Dang et al., 2024; Qiu et al., 2024). For instance, Dang et al. (2024) found a reduction in PV+ neurons in the BLA and their activation in rats eight weeks following a nerve injury caused by partial sciatic nerve ligation. Our results provide new evidence suggesting that neural plasticity involving PV^+^ neurons in the BLA plays an important role in mediating pain behavior following nerve injury.

By contrast, both excitation and inhibition of SST^+^ neuron in the BLA improved mechanical allodynia, suggesting a more nuanced role for these neurons in pain processing. The observation that both activation and inhibition of BLA SST^+^ interneurons produced similar pain-relieving effects in response to mechanical stimulation raises important questions regarding their circuit-level function. One potential explanation for this bidirectional effect is that SST^+^ neurons participate in homeostatic or compensatory mechanisms following nerve injury. Similar feedback loops have been observed in cortical circuits, where inhibitory plasticity adjusts to maintain network stability after injury or pathological hyperactivity (Harding and Salter, 2017; Dao et al., 2021). In the BLA, a comparable mechanism may exist, wherein any perturbation of SST^+^ neuron function—whether by increasing or decreasing their activity—disrupts maladaptive pain circuitry and restores nociceptive thresholds. This could be mediated by reciprocal connections between SST^+^ and other inhibitory interneurons, such as PV^+^ or vasoactive intestinal peptide (VIP^+^) neurons (Urban-Ciecko and Barth, 2016; Scheggia et al., 2020).

Interestingly, excitation of SST^+^ neurons in the BLA improved animal’s cold hypersensitivity but not their thermal hypersensitivity, while inhibition of SST^+^ neurons in the BLA does the opposite. This indicates that SST^+^ interneurons may play different role in circuits processing thermal and cold information. This aligns with findings in other brain regions where SST^+^ interneurons can either suppress excitatory output or disinhibit downstream targets depending on the state of the network (Cichon et al., 2017; Brockway et al., 2023). Our findings also raise a critical question whether pain processing across different modalities (mechanical, thermal, and cold) engages distinct neural circuits involving the same interneuron populations within the BLA. The fact that manipulation of BLA SST^+^ neurons affects pain behavior in all three modalities, while PV^+^ manipulation was only effective for mechanical and thermal hypersensitivities, suggests a potential link between nociceptive pathways and interneuron function. Cold hypersensitivity is often mediated by transient receptor potential (TRP) channels, including TRPM8 and TRPA1, which may preferentially interact with SST^+^-regulated circuits (Zhang et al., 2023). In contrast, mechanical and thermal pain pathways, which rely more heavily on TRPV1 and ASIC channels, may be more tightly coupled to PV^+^ interneuron function (Corder et al., 2019). Future studies employing cell-type-specific manipulations combined with nociceptive receptor tracing could further elucidate these relationships.

By dissecting the roles of PV^+^ and SST^+^ interneurons in the BLA, our study provides new insights into the circuit mechanisms underlying chronic pain. While PV^+^ excitation strongly modulates mechanical and thermal pain, SST^+^ neurons exhibit a more complex, bidirectional influence on nociceptive thresholds, particularly in cold allodynia. These findings suggest that distinct interneuron populations engage different nociceptive pathways and feedback loops, opening new possibilities for precision-targeted therapies for chronic pain. Beyond local inhibitory interneurons, our findings also implicate BLA-PFC and BLA-PAG circuits in chronic pain modulation. Previous studies have shown that inhibition of BLA projections to the PFC alleviates pain in mice with nerve injury (Gadotti et al., 2019), reinforcing the idea that amygdala-prefrontal pathways contribute to pain perception. Given that PV^+^ and SST^+^ neurons regulate BLA output, they may indirectly shape these descending pain-modulatory circuits, either amplifying or suppressing pain perception at a systems level (Shiers et al., 2018; Guo et al., 2022).

The results of the present study suggest several important directions for future research. First, high-resolution viral tracing and single-cell transcriptomics could help distinguish SST^+^ and PV^+^ subpopulations within the BLA, revealing whether distinct interneuron subtypes preferentially regulate specific pain modalities (Mihaljevic et al., 2019; Nagaeva et al., 2020). Second, targeted pharmacological strategies that enhance SST-peptide signaling or boost PV^+^ interneuron function could offer novel therapeutic avenues for chronic pain management (Harding and Salter, 2017; Brockway et al., 2023). Finally, investigating how neuromodulatory inputs (e.g., noradrenergic, cholinergic, etc.) interact with inhibitory microcircuits may clarify how these interneurons shape pain processing in a state-dependent manner (Liu et al., 2021; Rodenkirch et al., 2022; Slater et al., 2022; Sun et al., 2023). Understanding how these microcircuits involving PV^+^ and SST^+^ interneurons and their interaction with broader pain networks will be crucial for developing effective neuromodulation interventions for chronic pain conditions.

## Funding acknowledgement

This work was in part supported by NSF CBET 1847315 and NIH R01NS119813.

## Disclaimer of conflict of interest

Q.W. is the co-founder of Sharper Sense.

